# SHEN-LONG, a novel cardiac regulatory locus, regulates cardiac master transcription factor NKX2-5 in pluripotent stem cell derived cardiomyocytes

**DOI:** 10.1101/2024.12.24.630259

**Authors:** María Agustina Scarafía, Julia Halek, Joaquín Smucler, Ariel Waisman, Lucía Moro, Gustavo Sevlever, Santiago Miriuka, Alejandro La Greca

## Abstract

Cardiac development is a finely regulated process, transforming undifferentiated cells into the specialized cell components of the heart. Super-Enhancers (SE), clusters of enhancers densely populated with transcription factors, play a central role in cell fate decisions, including cardiogenesis. In this work, we studied a cardiac SE in chr3q25.31. This locus also contains a cardiac specific long non-coding RNA: LINC00881 (LINC881). We termed this region of interest SHEN-LONG (**S**uper **H**eart **EN**hancer and **LONG** non-coding LINC881), and selected it for further functional characterization. Our analysis revealed SHEN-LONG is a hotspot of cardiac kernel transcription factor binding sites. Cardiac differentiation of human iPSC with partial Knock Out of SHEN-LONG resulted in cardiomyocytes with reduced NKX2-5 expression, a master cardiac transcription factor, suggesting a significant role of SHEN-LONG in this process. Whole transcriptome sequencing of SHEN-LONG KO cardiomyocytes produced 134 differentially expressed genes located at great distances (*>*4Mb) emphasizing its long-range functional impact. A regulation mediated essentially by the SE and not by LINC881 was confirmed when LINC881 overexpression did not recover NKX2-5 expression. These findings provide insights into a novel NKX2-5 regulatory mechanism. The knowledge gained in this work may pave the way for advances in therapeutic interventions for cardiovascular disorders.

## 1 Introduction

Cardiac differentiation and development represent an intricate orchestration of molecular events that transform undifferentiated cells into the highly specialized and functional components of the heart [1]. The significance of understanding human cardiac development extends beyond its fundamental role in embryogenesis, and may provide crucial insights into the etiology of congenital heart diseases and facilitate the development of innovative therapeutic strategies.

Within the regulatory landscape of cardiac specification, the transcription factor NKX2-5 takes center stage as one of the cardiac kernel components, along with TBX5, GATA4, HAND1/2 and MEF2C [2]. Mutations in NKX2-5 or in its regulatory sequences (promoter or enhancers) have been associated with septal defects [3, 4], Tetralogy of Fallot [5], and issues in the development of the conduction system [6]. Moreover, complete or heterozygous deletion of NKX2-5 results in an embryonic lethal phenotype, that presents the absence of conduction system structures [7], and a decrease in its expression has serious consequences on normal cardiac functions [4, 8]. Nevertheless, in KO animal models not all cardiac structures are absent and *in vitro* cardiomyocytes with an altered genetic program have been obtained from NKX2-5-KO stem cells [9]. This indicates that NKX2-5 is not essential for the development of all cardiac structures, but it is crucial for overall heart function and development. NKX2-5 plays a pivotal role in directing the expression of genes critical for cardiomyocyte specification early in cardiac differentiation and repressing cardiac progenitor genes at later stages. Previous studies have identified SMAD proteins, YY1 and GATA4 among the regulatory elements of NKX2-5 activation [9, 10], but many aspects of its regulation are still unexplored.

Among the epigenetic factors of cardiogenesis, Super-Enhancers (SE) emerge as key components of cardiac development, governing cell identity by finely adjusting the expression of key developmental genes [11]. SE are constituted by clusters of enhancer elements densely populated with cell type-specific transcription factors, emerging as key players in the control of cell fate decisions [12]. Not all enhancer elements within a SE are functionally equivalent and may present additive, redundant or independent functions, also presenting a hierarchy where one element may exert a more powerful effect than the others [13, 14, 15]. Studying SE function is a challenging task given that disruption of large genomic regions can result in disturbance of genes involved in complex regulatory and transcriptional programs, leading to difficulties in the readout of specific biological effects. The most efficient strategy to study SEs is the genetic dissection of its individual constituent elements [16]. In contrast to classic enhancers, SE elements can function in an oriented-dependent manner [15] and may engage in strong interactions over large genomic distances [17]. Furthermore, SEs have been found to interact with long non-coding RNAs (lncRNAs)[18], adding a nuanced layer to this regulatory landscape. Together, SE and lncRNAs may exert their functions in mechanisms known as *in cis*, when regulation is exerted in close proximity and shared chromatin context, or *in trans* when their targets are localized over large genomic distances [19]. The intricate interplay of SEs and lncRNAs underscores their paramount roles in the complexities of cardiac differentiation.

In this work, we investigate the role of a previously reported but uncharacterized cardiac SE located in chr3q25.31. We found that this locus presents a hotspot of cardiac kernel transcription factor binding sites and thus represents an interesting element to study in more detail. Furthermore, *LINC00881* (LINC881), a primate and cardiac specific lncRNA, is transcribed from this genomic position. Recently, LINC881 has been shown to be regulated by NKX2-5 and participate in cardiac lipid droplet metabolism [20]. Here, we focused on the region comprising the binding site of cardiac kernel transcription factors in the SE and the first exon of LINC881, and named this region of interest SHEN-LONG (<u>**S**</u>uper <u>**H**</u>eart <u>**EN**</u>hancer and <u>**LONG**</u> non-coding RNA LINC881). Due to the primate-specific nature of this interesting region, we sought to investigate the role of SHEN-LONG in human cardiac differentiation based on the many advantages provided by human induced pluripotent stem cell (hiPSC)-cardiac differentiation model. Through loss and gain of function assays, we found that this SE is involved in the positive regulation of NKX2-5, and that this regulatory function is independent of LINC881 activity.

## 2 RESULTS

### SHEN-LONG comprises cardiac and primate specific elements: a Super-Enhancer and lncRNA LINC881

We identified a region of interest in human locus chr3q25.31 using publicly available data. The region is located between bases 157,070,000-157,150,000 and encompasses a cardiac SE that was reported to be active in cardiac tissue samples from left ventricle, right ventricle and right atrium (Figure 1A, top panel). This region is also the locus for three long non-codign RNAs (lncRNAs): *LINC02029, LINC00880* and *LINC00881* (LINC881), (Figure 1A, bottom panel). With respect to these genes, previous studies have documented a strong association of LINC881 with cardiac lineage [21], in agreement with tissue-specific gene expression databases that confirm a highly enriched expression of LINC881 in human cardiac tissues, but not in other tissues or organs (Figure 1B, data available at GTEx portal-release V8). Interestingly, *LINC02029* and *LINC00880* are not actively transcribed in cardiac tissue, despite sharing the locus with a cardiac SE, and in contrast, show enriched transcriptional activity in testis and pituitary gland respectively (Supplementary Figure 1A).

**Figure 1:**
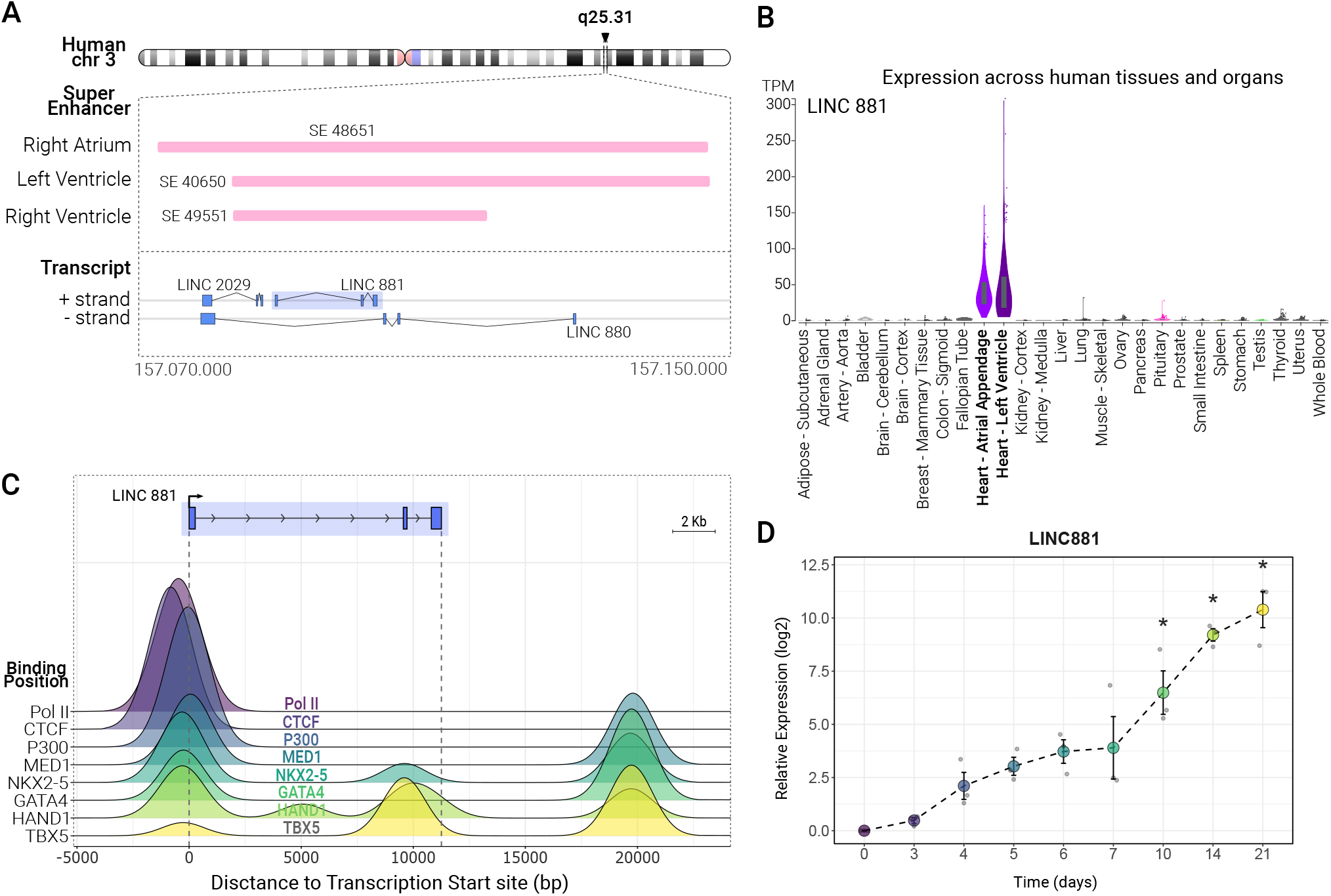
Characterization of genomic elements/features near SHEN-LONG. A) Super-Enhancer and lncRNA annotations in region chr3q25.31. Annotation of super enhancers SE 48651 (in the right atrium), SE 40650 (in the left ventricle) and SE 49551 (in right ventricle) was obtained from dbSUPER database. Transcript annotation of *LINC02029, LINC00881* (LINC881) and *LINC00880* was obtained from NCBI. B) LINC881 expression in human tissues and organs obtained from GTEx portal. C) Coverage analysis of binding sites of cardiac kernel transcription factors TBX5, HAND1, GATA4 and NKX2-5; transcription proteins PolII and p300; and enhancer related proteins MED1 (Mediator 1) and CTCF. ChIP-seq data was obtained from ChIP Atlas database D) Expression levels of LINC881 were assesed by RT-qPCR at nine time points of differentiation protocol. Statistical analysis was performed with one-way ANOVA followed by Tukey test. * = p*<*0.05.

We used public ChIP-seq data from pluripotent stem cell-derived cardiomyocytes available at the ChIP Atlas database [22] to study the binding profile of key cardiac transcription factors and other proteins near the locus of LINC881. Results revealed the presence of a cluster of binding sites of the cardiac proteins NKX2-5, GATA4, HAND1 and TBX5, enhancer related proteins MED1 and CTCF, and promoter proteins Pol II and p300 within *±* 1Kb of LINC881 transcription start site (Figure 1C).

To test the expression dynamics of LINC881 during cardiac differentiation, we differentiated human induced pluripotent stem cells (hiPSC) to cardiac lineage using a standardized protocol [23] and evaluated LINC881 expression by RT-qPCR at days 0, 3, 4, 5, 6, 7, 10, 14 and 21 (protocol design in Supplementary Figure 1D). We found that LINC881 expression increased gradually from day 0 up to day 21 (Figure 1D). As reported by previous studies, sequence alignment with BLAST tool confirmed that both the transcript of LINC881 and the genomic sequence of the putative promoter proved to be conserved in primates, but not in other species (Supplementary Figures 1B and C).

The reported elements comprised in locus chr3q25.31 show a specificity to cardiac lineage. Since the SE and the lncRNA LINC881 share the same genomic locus, there lied the possibility that they could play a role in cardiac differentiation by interacting with each other. In light of all the evidence of cardiac specific activity, and because SEs commonly participate in cell fate specification [12], we selected the region comprising part of the underlying cardiac SE, the first exon of cardiac lncRNA LINC881 and the hotspot of cardiac and regulatory protein binding sites to further characterize its role in cardiac differentiation. We named this region SHENLONG (**S**uper **H**eart **EN**hancer and **LONG** non-coding RNA LINC881).

### Knock Out of SHEN-LONG significantly alters NKX2-5 expression in cardiomyocytes

To better understand the role of SHEN-LONG in cardiac differentiation, we knocked out a fragment of 2,2 Kb (chr3:157,089,505-157,091,719) using CRISPR/Cas9 system (Figure S2A). The deleted region included various genomic elements: the binding site of cardiac transcription factors, the first exon of LINC881, and part of the underlying SE. With this strategy, we generated two knock out (KO) hiPSC lines: KO1 and KO2 (Figure 2A, top panel). Sanger sequencing confirmed deletion of SHEN-LONG in both cell lines (Figure S2B). We tested the expression of pluripotency core transcription factors NANOG, SOX2 and OCT4 by immunofluorescence and found no difference between SHEN-LONG KO cell lines and wild type (WT) hiPSC (Figure S2c), indicating that knock out of SHEN-LONG did not alter pluripotency markers.

**Figure 2:**
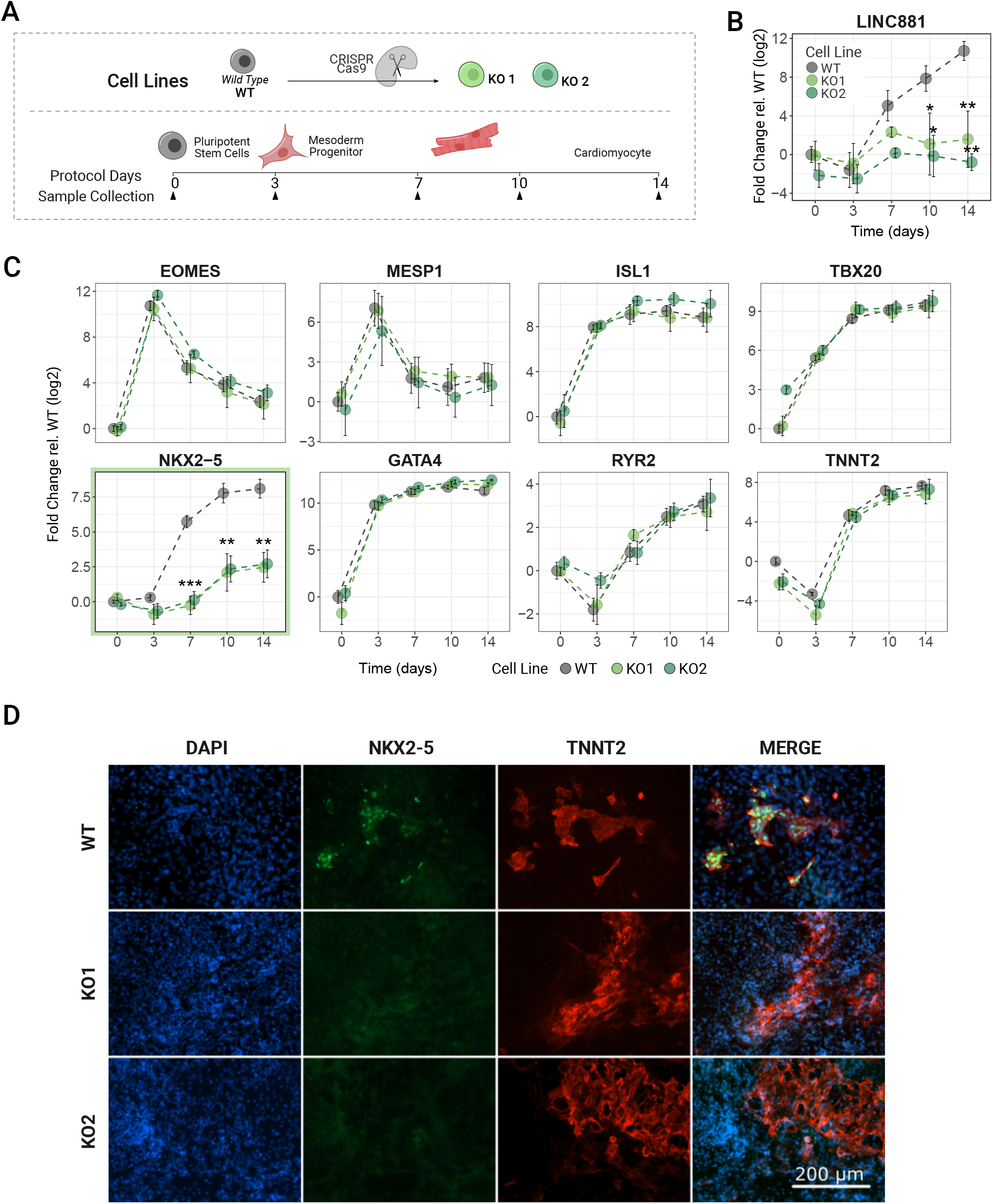
Cardiac differentiation of two SHEN-LONG Knock Out lines compared to Wild Type. A) Schematic of experimental design. Arrows show the days of RNA sample collection. B) Expression levels of LINC881 measured by RT-qPCR. C) Expression levels of gene markers: mesoderm commitment (*EOMES* and *MESP1*); cardiac progenitor transcription factors (*GATA4, ISL1, NKX2-5, TBX20*); cardiac ion channel (*RYR2*) and sarcomeric (*TNNT2*) specific protein genes. Data is presented as mean of three biological replicates with bars as mean *±* s.e.m. Statistical analysis was performed with one-way ANOVA followed by Tukey test. * = p*<*0.05, ** = p*<*0.01, *** = p*<*0.001. D) Immunofluorescence staining of NKX2-5 (green) and TNNT2 (red) at day 14 of cardiac differentiation in WT, KO1 and KO2 cell lines. Nuclei were stained with DAPI (blue). Merged channels are shown in the last column.

Next, we differentiated both KO and WT cell lines to cardiomyocytes. At day 14, differentiated cells from all cell lines showed spontaneous contractile activity, one of the hallmark characteristics of cardiomyocyes, indicating that the absence of SHEN-LONG did not prevent differentiation to cardiac lineage. We evaluated LINC881 expression at days 0, 3, 7, 10 and 14 by RT-qPCR and found that it was significantly downregulated in both KO lines compared to WT cell line at days 10 (KO1 p=0.045; KO2 p=0.026) and 14 (KO1 p=0.0022; KO2 p=0.00049) of the differentiation protocol (Figure 2B). These results validated the KO strategy employed.

To further study cardiac differentiation of SHEN-LONG KO cell lines, we evaluated the expression levels of several cardiac differentiation markers at days 0, 3, 7, 10 and 14 by RT-qPCR (Figure 2C). Of the 8 genes evaluated, 7 showed no differences between KO and WT lines, including gene markers related to mesoderm formation (*EOMES, MESP1*); cardiac progenitor transcription factors (*ISL1, TBX20, GATA4*); and cardiomyocyte self-proteins (*TNNT2, RYR2*). This supported the idea that cardiac differentiation was not impeded due to SHEN-LONG deletion. However, the expression of the cardiac master transcription factor *NKX2-5* was significantly downregulated in both KO lines compared to WT on day 7 (KO1 p=0.00014; KO2 p=0.00021), day 10 (KO1 p=0.00744; KO2 p=0.00932) and day 14 (KO1 p=0.00411; KO2 p=0.00522).

To assess whether the decrease in *NKX2-5* mRNA was accompanied by an alteration in the protein levels, we examined the presence of NKX2-5 protein in KO cardiomyocytes by immunofluorescence (Figure 2D). Neither KO1 nor KO2 cell lines presented NKX2-5^+^ cardiomyocytes (TNNT2^+^ cells) at day 14 of differentiation, indicating that *NKX2-5* expression was altered both at transcript and protein level. These striking results show that SHEN-LONG-KO hiPSC were able to differentiate to beating cells with an altered expression of *NKX2-5*, a central component of cardiac kernel that is conserved from *D. melanogaster* to mammals.

In summary, these results indicated that, while cardiac differentiation was not prevented, the cardiac gene program was altered in SHEN-LONG-KO cardiomyocytes.

### Knock out of SHEN LONG results in an altered expression of genes at great distances

We next sought to identify the repertoire of genes under the regulation of SHEN-LONG by whole transcriptome sequencing. Since both SHEN-LONG KO lines, KO1 and KO2, showed similar expression levels of cardiac differentiation marker genes, we selected only KO1 for sequencing experiments. Next, we performed total RNA sequencing on KO1 and WT cardiomyocytes on days 7 and 10 of the differentiation protocol. We identified a total of 134 differentially expressed (DE) genes between cell lines: 81 DE genes at day 7 and 101 DE genes at day 10 (Figure 3A). Interestingly, we found that most DE genes are downregulated in the KO condition, while approximately 6% and 15% of genes were upregulated in KO cardiomyocytes on day 7 and 10 respectively. In agreement with our qRT-PCR results, *NKX2-5* was found to be significantly downregulated in the KO condition. Also, fold change of the expression levels was greater in down-regulated genes compared to upregulated genes. This could be indicating that the role of SHEN-LONG would be related to a positive regulation of its target genes.

**Figure 3:**
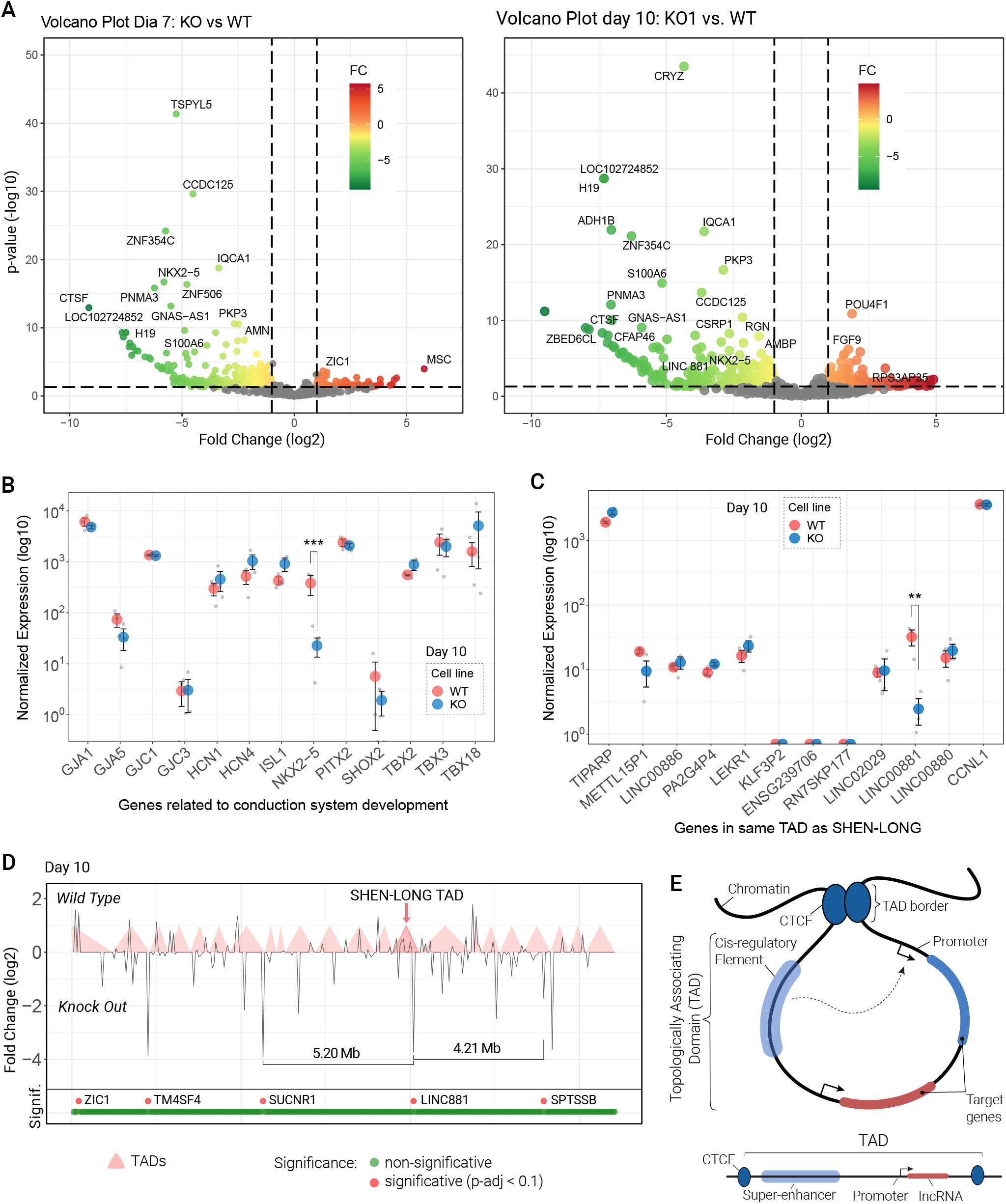
Differential expression analysis of SHEN-LONG KO and WT day 10 cardiomyocytes. A) Volcano plot representation of expressed genes in WT and SHEN-LONG KO cardiomyocytes at day 7 (left) and day 10 (right). Differentially expressed genes are shown colored by Fold Change (log2), non differentially expressed genes are shown in gray. B) Normalized expression levels in log10 scale of genes related to conduction system development. *** = p-adj*<*0.001 compared between KO (red) and WT (blue) cardiomyocytes at day 10. C) Normalized expression in log10 scale of genes localized in TAD 749 (same as SHEN-LONG) between KO (red) and WT (blue) cardiomyocytes at day 10. ** = p-adj*<*0.01. D) Relative localization, TAD and distance of diferentially expressed genes located within 10 Mb up and downstream of SHEN-LONG (distances not to scale) at day 10 cardiomyocytes. In total 299 genes are shown. Statistical significance is shown in the bottom panel using p-adj*<*0.1 as significance threshold. The distence between the two differentially expressed genes closest to SHEN-LONG are pointed out: SUCNR1 at 5.20 Mb, and SPTSSB at 4.12 Mb. TADs are shown as pale red rectangles. E) Schematic of basic structure and elements of a Topologically Associating Domain (TAD).

Since alterations in *NKX2-5* expression have been associated to defects in cardiac conduction system development, we evaluated the expression levels of various genes related to its development, including several connexins (*GJA1, GJA5, GJC1, GJC3*), potassium channels (*HCN1, HCN4*) and transctiption factors (*ISL1, NKX2-5, PITX2, SHOX2, TBX2, TBX3, TBX18*). With the exeption of *NKX2-5*, we found that none of these genes were differentially expressed at days 7 (Figure S3A) or 10 (Figure 3B) in our model indicating that, at these timepoints, there would be no evidence of conduction system compromise.

Then we evaluated the possibility that SHEN-LONG exerted its regulatory function within local chromatin organization on genes in the proximity of its locus (in *cis*) or at distant genomic loci (in *trans*). To this end, we analyzed the expression levels of genes localized in the same Topologically Associating Domain (TAD). Of the 12 genes present in the same TAD, only LINC881 showed differential expression between WT and KO cardiomyocytes, but none of the remaining genes were affected by SHEN-LONG KO at day 10 of differentiation protocol (Figure 3C) or at day 7 (Figure S3B). When analyzing gene expression 10 Mb upstream and downstream of SHEN-LONG, a region comprising 299 genes, we found the two nearest DE genes to be located at 5.20 Mb upstream (7 TADs away) and at 4.21 Mb downstream (5 TADs away) at day 10 (Figure 3D). These represent greater distances than tipically reported in *cis* regulatory mechanisms. Similar results were seen at day 7 (Figure S3C). We also found no grouping of DE genes in any particular location in the human genome (Figure S3D).

These results are not consistent with a *cis* regulatory mechanism and together they support that SHEN-LONG would be exerting its function in *trans*, over large genomic distances.

### NKX2-5 expression levels are not restored after LINC881 Over Expression

With the aim of understanding if the regulation of *NKX2-5* expression by SHEN-LONG was dependent on both elements contained in the genomic region deleted in the KO lines, we evaluated if LINC881 overexpression could restore *NKX2-5* levels in SHEN-LONG KO cardiomyocytes. To this end, we cloned LINC881 cDNA into an expression vector under a doxycycline promoter and generated LINC881 overexpression cell lines from both KO cell lines, rescue lines R1 and R2 respectively (Figure 4A, top panel, and Figure S4A). First, we corroborated that doxycycline had no negative effect in cardiac differentiation using the WT stem cell line, treating cells with or without doxycycline between days 3 and 10, and assessing expression levels of cardiac differentiation markers: *EOMES, MESP1, ISL1, TBX5, TBX20, GATA4, RYR2, TNNT2, NKX2-5* and LINC881 (Figure S4B). There was no detectable difference between treatments and all markers showed similar expression levels to standard condition.

**Figure 4:**
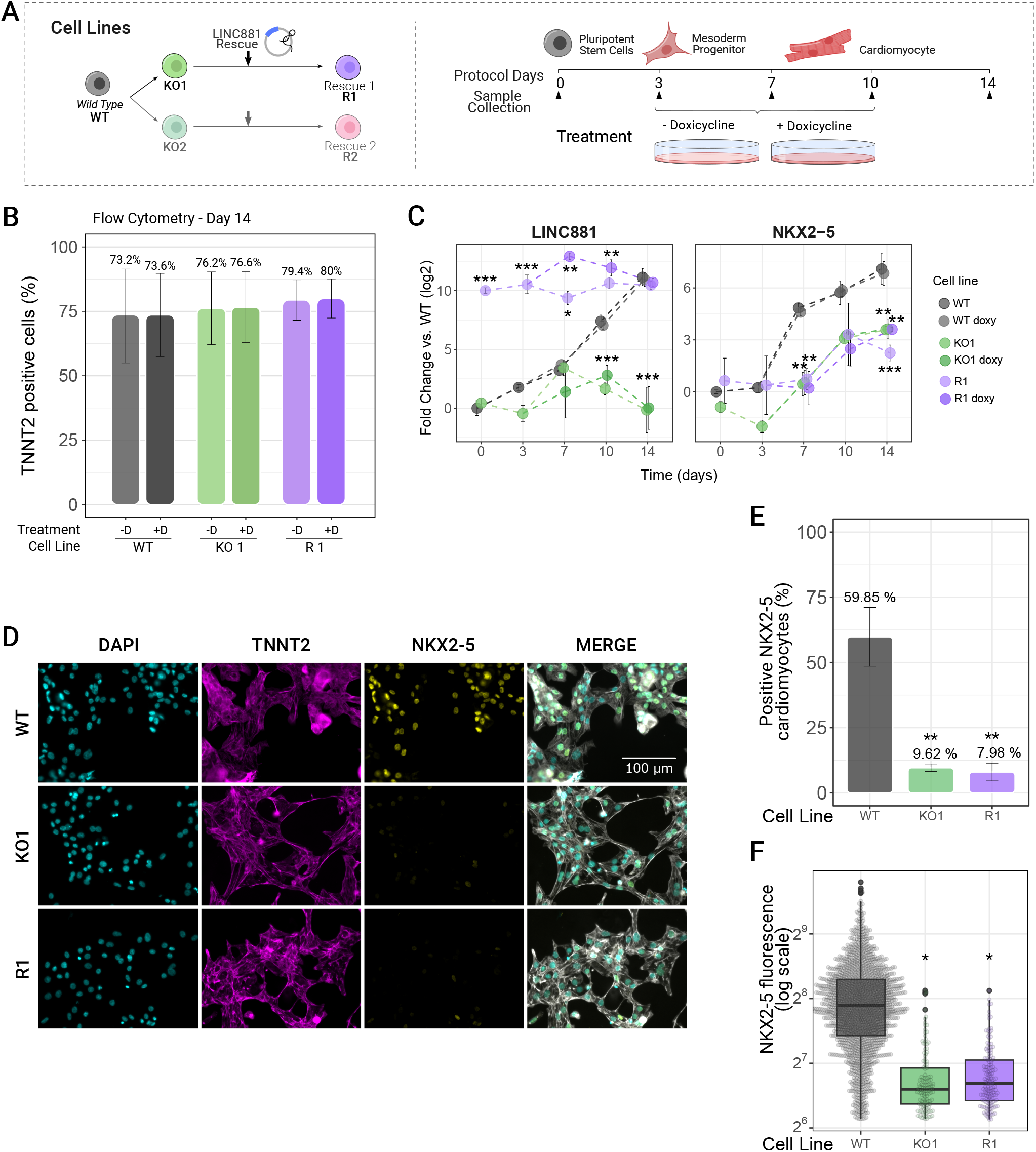
Cardiac differentiation of LINC881 in WT, KO1 and R1 cell lines. A) Schematic of differentiation protocol (left) and cell lines used (right). B) Cardiomyocyte percentages at day 14 measured as percentage of TNNT2^+^ cells, with and without doxycycline treatment. C) Expression levels of LINC881 and NKX2-5 along cardiac differentiation protocol, with and without doxycycline treatment assesed by RT-qPCR in WT, KO1 and R1 cell lines. Data is presented as mean of three biological replicates with bars as mean *±* s.e.m. Statistical analysis was performed with one-way ANOVA followed by Tukey test or Kruskal-Wallis followed by Dunn-Bonferroni when assumptions were not met. Statistical significance is reported relative to WT cell line values. * = p*<*0.05, ** = p*<*0.01, *** = p*<*0.001. D) Immunofluorescence staining of NKX2-5 (yellow) and TNNT2 (magenta) at day 14 of cardiac differentiation in WT, KO1 and R1 cell lines. Nuclei were stained with DAPI (cyan). Merged channels are shown in the last column, with TNNT2 in gray. E) Representative quantification of NKX2-5 mean fluorescence per nuclei in cardiomyocytes (TNNT2^+^/NKX2-5^+^ cells) for WT, KO1 and R1 cell lines. Boxplots show the quartiles, with the median as athick black line. Statistical significance was performed with one-way ANOVA of the median of three biological replicates followed by Tukey test: * = p*<*0.05.

We then differentiated all five cell lines (WT, KO1, R1, KO2 and R2) into cardiomyocytes using the protocol described before, treating cells with or without doxycycline between days 3 and 10 (Figure 4A, right panel). All cell lines produced similar percentages of cardiomyocytes at day 14 (mean *≈* 73-80%) measured as TNNT2^+^ cells by flow cytometry (Figures 4B and S4C). All cell lines showed spontaneous contractile activity. This indicated that neither the overexpression of LINC881 nor the doxycycline treatment altered the ability to differentiate into cardiomyocytes or affected contractile activity.

Next, we evaluated LINC881 and *NKX2-5* expression by RT-qPCR in WT, KO1 and R1 cells (Figure 4C). We found that LINC881 expression was low in KO1 cell line (green) throughout the diffentiation protocol compared to WT (gray), showing statistical significance at days 10 and 14 (p*<*0.001). Expression levels were similar with and without doxycycline treatment. On the other hand, R1 cells (violet) showed an elevated expression of LINC881 that was significantly higher than WT at days 0 (p= 0.00001), 3 (p= 0.00016), 7 (p= 0.0389 doxy-/0.0059 doxy+) and 10 (p= 0.01972 doxy-/0.00155 doxy+), but showing no diferences at day 14 compared to WT cardiomyocytes. Importantly, *NKX2-5* expression levels were similar between KO1 and R1 cell lines, and significantly downregulated compared to WT cells on day 7 (KO1 p=0.00351; KO1 doxy p=0.00416; R1 p=0.00593; R1 doxy p=0.00231) and 14 (KO1 p=0.00749; KO1 doxy p=0.00822; R1 p=0.00051; R1 doxy p=0.00782). Similar results were obtained in KO2 and R2 cell lines (Figure S4D). This is consistent with the analysis of TNNT2 and NKX2-5 positive cells on day 14 through flow cytometry where we observed that KO1 and R1 cell lines presented no NKX2-5^+^ cardiomyocyes (TNNT2^+^ cells) (Figure S5A), in contrast with the WT cells where approximately half of cardomyocytes obtained were NKX2-5^+^. Since elevated levels of LINC881 in R cell lines were comparable to levels found at day 10/14 WT cardiomyocytes, even in the absence of doxycycline treatment, we decided to perform further experiments without adding the doxycycline.

Our previous experiments to study NKX2-5 protein expression by immunofluorescence had been done directly over the intact monolayer culture at day 14 of the cardiac protocol. In this context, the assembly of many three-dimensional structures complicates the proper visualization of individual cells and nuclei. To surpass this limitation and better study NKX2-5 protein expression in the rescue model, we performed the selection of cardiomyocytes by implementing a Selection Medium and reseeding the cells in less confluency prior to the immunostaining analysis (See Methods section). After cardiomyocyte selection, immunofluorescence of TNNT2 and NKX2-5 was performed at day 22. In all cell lines, a high proportion of cardiomyocytes can be observed as TNNT2^+^ cells, with very few nuclei found in TNNT2^*−*^ cells (Figure 4D). In WT cardiomyocytes, a high proportion of nuclei exhibit strong fluorescence for NKX2-5, whereas in KO1 and R1 cell lines, fewer nuclei are NKX2-5^+^, and the fluorescence appears dimmer. Nuclei were identified with DAPI staining, and mean fluorescence per nuclei was quantified for NKX2-5 staining, distinguishing between TNNT2 positive and negative cells. In line with the RT-qPCR results, a lower percentage of NKX2-5^+^ cardiomyocytes was found in KO1 and R1 cell lines, compared to WT NKX2-5^+^ cardiomyocytes (9.62% and 7.98% vs 59.85%, Figures 4E and S5B). We also found that among NKX2-5^+^ cardiomyocytes, KO1 and R1 presented a significantly reduced expression of NKX2-5, measured as lower mean fluorescence. Population median: WT 328.7 *±* 100.4; KO1 137.8 *±* 46.7; R1 136.8 *±* 34.4 (Figure 4F). Similar results were obtained for KO2 and R2 cell lines (Figures S4E, S4F and S4G. Percentage of NKX2-5^+^ cardiomyocytes: WT 55.3%, KO2 7.9% and R2 8.8%. Population median: WT 318.0 *±* 88.3; KO1 140.2 *±* 38.9; R1 140.2 *±* 63.9).

Altoghether theses results indicate that while the locus of SHEN-LONG regulates the expression of *NKX2-5*, the RNA transcript of LINC881 would not be sufficient to exert this regulatory role on its own, and on the other hand, it suggests that the enhancer element in the genomic loci would be necessary for this regulatory mechanism.

## 3 Discussion

Our results show that SHEN-LONG, a primate-specific locus previously uncharacterized, participates in the regulation of NKX2-5 during human cardiac differentiation. To our knowledge, this is the first report functionally connecting NKX2-5 expression to SHEN-LONG. This locus is located in chromosome 3 and comprises primate and cardiac specific elements: the long non-coding RNA *LINC00881* and a Super-Enhancer that contains a hotspot for cardiac transcription factor binding sites. To explore the function of SHEN-LONG locus in cardiac differentiation, we performed loss of function assays by generating KO cell lines from human iPSC. Cardiomyocytes derived from KO cell lines presented a significant down-regulation of cardiac master transcription factor NKX2-5, both at the transcript and the protein levels, but not in other cardiac transcription factors, evidencing a perturbation in the normal cardiac gene program, and revealing a new element involved in NKX2-5 regulation.

The results presented here go in line with the work of Anderson *et al* ([9]) where they produced contractile human cardiomyocytes in the absence of NKX2-5 (NKX2-5-KO). Despite presenting normal expression levels in most cardiac transcription factors, the authors reported an altered cardiac gene program featuring the persistent expression of early cardiac progenitor marker PDGFRa at day 42, and the diminished expression of VCAM1 along the differentiation protocol, both direct targets of NKX2-5. The authors concluded that the resulting phenotype would imply an incomplete differentiation of NKX2-5 -/- cardiomyocytes, with an immature gene program. Similarly, in our study, SHEN-LONG-KO cell lines differentiated to contractile cardiomyocytes despite presenting a substantial downregulation of NKX2-5. Furthermore, expression levels of cardiac transcription factors TBX5, TBX20, GATA4 and ISL1 did not show differences between WT and KO cell lines assesed by qPCR (at days 3, 7, 10 and 14), and no cardiac master transcription factors were found differentially expressed in RNA-seq analisys at the evaluated timepoints (days 7 and 10). It remains to be studied whether the downregulation of NKX2-5 is sustained at later stages of cardiac differentiation, and if the phenotype of our KO-derived cardiomyocytes ressembles the one reported by Anderson *et al*.

Although our results demonstrated the involvement of SHEN-LONG in NKX2-5 regulation, we had no clarity on the level of participation and interactions of its two inner elements. To address this point, we sought to perform a gain-of-function assay to make an independent assessment of the function of LINC881 and the Super-Enhancer by restoring LINC881 expression in KO cell lines. First, to understand the type of regulatory mechanism exerted by SHEN-LONG, we performed RNA-seq on KO1 and WT-derived cardiomyocytes. The analysis revealed that the nearest up-and downregulated genes were found in TADs several Kb away, providing evidence of a trans-orchestrated mechanism. In addition, differentially expressed genes were not grouped in any particular region of the genome, further supporting this notion. These results provided the basis for the overexpression strategy: a mechanism in trans would imply that the function of LINC881 is not dependent on the chromatinic context in which it is transcribed, while a mechanism in cis would require the restoration of LINC881 expression in the same original biological or chromatinic context ([24]). Thus, the restoration of LINC881 expression in a heterologous manner was a sound and reasonable approach to discriminate between the two.

NKX2-5 downregulation in SHEN-LONG-KO cardiomyocytes highlighted the role of the locus in cardiac differentiation. With this result, we contemplated three possible scenarios with respect to the function of SHEN-LONG’s inner elements: first, that LINC881 transcript was sufficient for NKX2-5 expression, and the Super-Enhancer was not necessary; second, that the SuperEnhancer was sufficient for NKX2-5 expression, but LINC881 transcript was not necessary; third, that both elements were necessary for the expression of NKX2-5 due to an interaction between the two. However, the overexpression of LINC881 in both R cell lines, did not restore NKX2-5 expression levels in SHEN-LONG-KO cardiomyocytes. In the R1 and R2 cell lines, the transcript of LINC881 is present in the absence of the genomic region that corresponds to the Super-Enhancer. Therefore, the reduced expression of NKX2-5 in both R lines highlights the role of the genomic region as essential for NKX2-5 regulation.

Taking our results into account, we were able to conclude that two scenarios out of the three are possible and exclude the possibility that LINC881 on its own is sufficient to regulate NKX2-5. In contraposition, the evidence indicates that the Super-Enhancer is necessary for the positive regulation of NKX2-5. The two remaining possibilities are that both LINC881 and the Super-enhancer interact with each other to exert this regulatory role together, on one hand, and that the Super-Enhancer on its own is sufficient to regulate NKX2-5, on the other.

The results we obtained remain connected to and complement the findings in the study of Han *et al*. 2023 ([20]) that identified NKX2-5 as a positive regulator of LINC881 expression. In their work, they performed the KO of LINC881 transcript from the transcription start site to transcription termination site. LINC881 -/-cardiomy-ocytes presented an alteration in lipid droplet function at day 40 of cardiac differentiation, but authors reported NKX2-5 expression levels were not affected by loss of LINC881. Inversely, the downregulation of NKX2-5 produced a significant reduction in LINC881 expression levels. In light of these results, it is reasonable to hypothesize that the regulation of NKX2-5 in our model is being exerted by the genomic region corresponding to the Super-Enhancer alone. However, further investigation is needed to confirm this and dissect the underlying molecular mechanism.

In regard to this last finding, SEs consist of inner elements that present complex interactions. Here, we were able to establish an important function of SHEN-LONG in cardiac differentiation. However, we did not explore in depth the sub-elements of this Super-Enhancer. It may be possible that more than one sub-element is present within the 2.2 Kilobases knocked out in our model. Dissecting the inner elements of this super enhancer may shed light into SHEN-LONG’s nature and allow the distinction of sub-elements or facilitators ([16]) and further clarify the relevance of SHEN-LONG in cardiac differentiation and homeostasis.

Alongside these results, it has been established that NKX2-5 presents an evolutionary conserved autoregulation [25]. One evidence sustaining this argument is the presence of an NKX2-5 binding site in its own promoter. Also, in vertebrates, NKX2-5 presents a cooperative transcriptional regulation with other proteins, such as MEF2C and GATA4, by which NKX2-5 indirectly self-regulates [25, 10, 26]. ChIP-seq data showed that NKX2-5 also binds to SHEN-LONG, though the outcome of this binding has a function that is still unkown. It has been shown that NKX2-5 -/-cariomyocytes present a downregulation of LINC881 transcript, but it remains unclear whether it also produces the absence or reduction in SHEN-LONG activity. Taking the sum of our results into consideration, it is possible to assume that there are more elements participating in NKX2-5 indirect autoregulatory mechanism, that instead, could be dependent on other elements, such as distant enhancers or transcription factors, and could closely resemble a positive feedback loop or cooperative regulation with SHEN-LONG enhancer.

Hopefully, our results could serve as a focal point not only for understanding the complexities of heart development, but also in the pursuit of regenerative medicine strategies, and the potential translation of this knowledge into novel therapeutic interventions for cardiovascular disorders.

## 4 METHODS

### iPSC culture and cell lines

Human induced pluripotent stem cell (hiPSC) line FN2.1 (WT) was previously developed in Miriuka Lab from foreskin fibroblasts [27] and registeded in the cell line registry platform hPSCreg (https://hpscreg.eu/) under the name INEUi002-A. FN2.1-WT derived cell lines are registered as INEUi002-A-3 (KO1), INEUi002-A-4 (KO2), INEUi002-A-5 (R1) and INEUi002-A-6 (R2). Normal male karyotype was confirmed and their identity was authenticated by Short Tandem Repeats (STR). Pluripotency was asessed by expression of pluripotency core transcription factor genes by immunofluorescence (Figures S2C and D).

Cells were maintained in feeder-free conditions on plates coated with Geltrex (ref. A1413302, Thermo Fisher Scientific) in StemFlex defined medium (ref. A3349401, Thermo Fisher Scientific), replacing it each day. Cells were passaged every 3-4 days to maintain a confluence under 60-70% using Versene (ref. 15040066) following manufacturers instructions. Cell culture medium was supplemented with 10 *µ*M Rho kinase inhibitor (ROCKi) Y-27632 (ref 1254, Tocris) the first 24h.

### Cardiac differentiation

For cardiac differentiation, the protocol deveopled by Lian *et al*. was used with minor modifications [23]. A schematic of the protocol is shown in Supplementary figure 1D. Briefly, cells were dissociated into single cells with TrypLE Select 1X (ref. A1217702, Thermo Fisher Scientific) and seeded at 200-300,000 cells/24 well on Geltrex-coated 24-well plates in StemFlex defined medium + 10*µ*M ROCKi Y-27632 for 24h. Cells were maintained for three days until reaching 100% confluency. Then, cells were treated with CHIR99021 (ref. 4423, Tocris) for 24h at a final concentration of 10*µ*M in RPMI1640 medium (ref. 22400-089, Thermo Fisher Scientific) supplemented with B27 without insulin (ref. A1895601, Thermo Fisher Scientific). Untreated pluripotent cells were collected as day zero of the protocol. On day 3, differentiating cells were treated with IWP2 (ref. 3533, Tocris Bioscience) at a final concentration of 5*µ*M for 48h. Untreated cells were collected as day 3 of the protocol. At day 7 media was renewed. From day 9 onwards, media was replaced every two days with Basal Medium: RPMI1640 supplemented with B27 with insulin (ref. 17504-001, Thermo Fisher Scientific). Cardiac differentiation protocol was continued until day 14 or 21, when final sample was collected.

Alternatively, for quantitative immunofluorescence assays, cardiomyocytes were first selected. Briefly, at day 11 of differentiation protocol, cells destined to immunofluorescence analysis were washed with 1X PBS and incubated with 1X Trypsin-EDTA (ref. 25200-072, Thermo Fisher Scientific) for 5 minutes. After incubation, cell were resuspended in RPMI1640 supplemented with Fetal Bovine Serum 20% and centrifuged at 300 xg during 5 minutes. Cell pellet was resuspended in Replate Medium: RPMI1640 supplemented with B27, 10*µ*M Y-27632 and 10% KSR (ref. 10828028, Thermo Fisher Scirentific) and seeded on Geltrex-coated 6-well plates at a ratio of 1:1 (24 well:6 well). At day 12, culture media was replaced with Selection Media: RPMI1640 without glucose (ref. 11879020, Thermo Fisher Scientific) supplemented with 7mM sodium DL-lactate (ref. L4263, Sigma), 213 *µ*g/ml L-ascorbic acid 2-phosphate (ref. A8960, Merck) and 0.5 *µ*g/ml BSA fraction V (ref. 15260, Thermo Fisher Scientific). Selection media was renewed every 2 days until day 18. At day 18 media was replaced with Basal Media. On day 21, cells were detached as described before, and plated on circular glass coverslips previously treated 1 hour with Geltrex 1X in Replate Medium. The next day, cells were fixed in the glass coverslips.

### RNA isolation and RT-qPCR for transcript expression assessment

Total RNA was prepared with TRIzol Reagent (ref. 15596018, Ambion) following manufacturer’s instructions, treated with DNAse I (ref. 18047-019, Thermo Fisher Scientific), and then reverse transcribed into cDNA using MMLV reverse transcriptase (ref. M1705, Promega) and random primers (ref. 48190011, Invitrogen). Quantitative real time PCR (qPCR) was performed in a StepOne Real Time PCR system (Applied Biosystems) using FastStart Universal SYBR Green Master (Rox) (ref. 4913914001, Roche). Gene expression was quantified, normalized to the geometrical mean of HPRT1 and RPL7 housekeeping genes, data was then log2 transformed and relaivized to the average of the biological replicates for day 0 Wild Type condition. Values are presented as the mean *±* standard error of the mean (s.e.m.). Statistical significance for qPCR results was analyzed by one-way ANOVA with randomized block design followed by Tukey’s multiple comparisson test. When ANOVA’s assumptions were not met, data was analyzed with Kruskal-Wallis test followed by Dunn-Bonferroni correction. Primer sequences are detailed in Supplementary Table 1.

### Immunofluorescence staining

For immunofluorescence assay of cardiomyocytes at day 14, upon reaching day 14 of cardiac differentiation cells were fixed in 24 well plates, without modifying the protocol. For quantitative immunofluorescence assay, selected cardiomyocytes at day 22 were fixed in glass civerslips.

For pluripotent stem cells: cells were dissociated into single cells with TrypLE Select 1X and seeded at 25.000 cells/glass coverslip (1 cm^2^) in StemFlex medium supplemented with 10*µ*M ROCKi for 24 hs. Culture media was changed every day for 2-3 days.

Upon reaching necessary conditions, cells were washed three times with 1X PBS and then fixed for 10 minutes with 4% PFA at room temperature (RT). Cells were then permeabilized and blocked for 30 minutes with Permeabilization Buffer (PBSTS: 1X PBS, 0.1% Triton, 3% Normal Goat Serum). Samples were resuspended and incubated with Primary antibody dilution (1:50-200 depending on manufacturer recomendations in PBSTS) for 30 minutes at RT. Samples were washed with Wash Buffer (WB: 1X PBS, 0.1% Triton) three times and then incubated with Secondary antibody dilution (1:400 in PBSTS) for 1 hour at RT. Samples were washed with WB two times and then incubated with 0,5 *µ*g/ml DAPI (ref. D1306, Thermo Fisher Scientific) for 5 minutes. Then, samples were washed with WB 5 minutes and then rinsed with destiled water twice before mounting the coverslips on slides with 5*µ*l of Mowiol mounting media. Slides were left overnight at room temperature and coverslips were sealed the following day with clear nail polish. Samples were photographed on an inverted microscope EVOS XL Core Imaging System (Thermo Fisher Scientific). For quantification of day 22 cardiomyocytes, automatic nuclear segmentation was made using StarDist[28], and nuclear fluorescence intensity measurements for different channels was calculated with custom Fiji/ImageJ Macros. Measurement tables were then analyzed using custom R scripts with the tidyverse library. primary antibodies used are listed in Supplementary Table 2.

### Flow cytometry analysis

At day 14 hiPSC-derived Cardiomyocytes (hPSC-CM) were washed three times with 1X PBS and treated for up to 10 minutes with 10X TrypLE at 37ºC. The incubation was stopped with DMEM supplemented with 10% Fetal Bovine Serum (FBS). Cells were centrifuged at 300 xg 5 minutes, resuspended and then fixed for 10 minutes with 4% PFA at RT. Cells were permeabilized and blocked for 30 minutes with Permeabilization Buffer (PBSTS: 1X PBS, 0.1% Triton, 3% Normal Goat Serum). After centrifugation, samples were resuspended and incubated with Primary antibody dilution (1:50-200 depending on manufacturer recomendations in PBSTS) for 30 minutes at RT. Samples were washed with Wash Buffer (WB: 1X PBS, 0.1% Triton) and cetrifuged at 300 xg during 5 minutes three times. Cells were then resuspended and incubated with Secondary antibody dilution (1:400 in PBSTS) for 1 hour at RT. Samples were washed with WB and centrifuged at 300 xg during 5 minutes three times and resuspended in 1X PBS. Cell suspension was analyzed in C6 Accuri Cytometer, at least 10.000 events were counted at low speed for each condition. Primary antibodies used are listed in Supplementary Table 2.

### Generation of SHEN-LONG Knock Out cell lines

Two sgRNAs were designed using Benchling (benchling.com/). The oligonucleotides were annealed and cloned into the pSpCas9(BB)-2A-Puro v2.0 plasmid (ref. Addgene 62988) as described before [29]. The plasmids were cotransfected into iPSC cells with Lipofectamine Stem Reagent (ref. STEM00015, Thermo Fishser Scientific) in mTeSR media (ref. 100-0276, Stemcell Technologies) supplemented with 10*µ*M Y-27632 for 24 hs. Transfected cells were selected with 0.5*µ*g/ml puromycin for 48 hs and expanded in StemFlex media with 10*µ*M Y-27632 for 4-5 days. After selection, cells were dissociated with 1X TrypLE and individually picked up and transferred to 96-well plate in StemFlex media with 10*µ*M Y-27632 for clonal selection and expansion. Once colonies were obtained, cells corresponding to the same clone were passaged to two wells: one to generate stock and another for genomic DNA extraction. Deletion of region of interest was confirmed by end point PCR using primers TGTCGAGAGCACTGATTTCCC and CTGCTGCTACAGACTCCGAC.

### Whole transcriptome sequencing (RNA-seq)

Total RNA sequencing and data processing. TRIzol-purified samples were quantified with Qbit and 5*µ*g of total RNA was treated with DNase I and precipitated with 0.1 volumes of 3M NaAc pH=5.5 and 2 volumes of 100% ethanol. Library preparation and RNA sequencing was performed by Macrogen, South Korea. Libraries were built using TruSeq Stranded Total RNA with Ribo-Zero kits. Fastq files were aligned to hg38 genome with STAR software, and gene count was done with HTSeq. Differential expression analysis was done with DESeq2 package in R software (v4.1.2).

### Generarion of LINC881 Overexpression cell lines

For over expression (OE) of LINC881 to generate R cell lines, cDNA was synthesized from 1*µ*g RNA obtained from day 21 cardiomyocytes using 250ng Oligo(dT) Primer (ref. 18418-012, Invitrogen). LINC881 sequence was amplified trough PCR with primers AAACCGGTA-GAAAGTGCTAGGCACGTAGG and TTGCTAGCCTCA-CAATTCTGGAGGCCAGA, and cloned into plasmid vector ePB-HA-Oct6-P2A-mCherry developed in Miriuka Lab [30]. Plasmid was transfected into KO1 and KO2 cell lines, and cells were selected clonally as described before. Construct incorporation was confirmed by RT-qPCR of LINC881 transcript in pluripotent stem cell lines.

### Bioinformatic analysis and external data used

Cardiac Super Enhancer annotations were obtained from dbSUPER database [31] (asntech.org/dbsuper) and transformed to Hg38 with LiftOver tool. Binding site annotations for transcription factors NKX2-5, GATA4, HAND1 and TBX5, enhancer related proteins MED1 and CTCF, and promoter proteins Pol II and p300 were downloaded from ChIP-Atlas (https://chip-atlas.org/)[22] and GEO (accession numbers: GSE133833 and GSE89457). Bedtools (v2.30.0) intersect uas used to keep annotations in LINC881 locus and Bedtools coverage to calculate number of annotation in each base. Obtained data was imported and plotted in R.

Expression of long non-coding RNAs in adult human tissues was obtained from GTEx portal, V8.

Transcript sequence annotations were obtained from NCBI. Basic Local Alignment Tool (BLAST) was used with default parameters to asses conservation levels of LINC881 transcript and promoter sequence in *Mammalia* class (taxid 40674). Results were analized and plotted in R. Only results showing a length conservation greater than 50% were analyzed, or until a genus outside primates appeared.

Topollogically Associating Domains (TADs) coordinates were obtained from Encode Project (https://www.encodeproject.org/), and transformed to Hg38 assembly using liftOver tool.

## Data availability

The generated data supporting the findings of this study will be available upon publication

## SUPPLEMENTARY INFORMATION

### Supplementary Tables

**Table 1.**
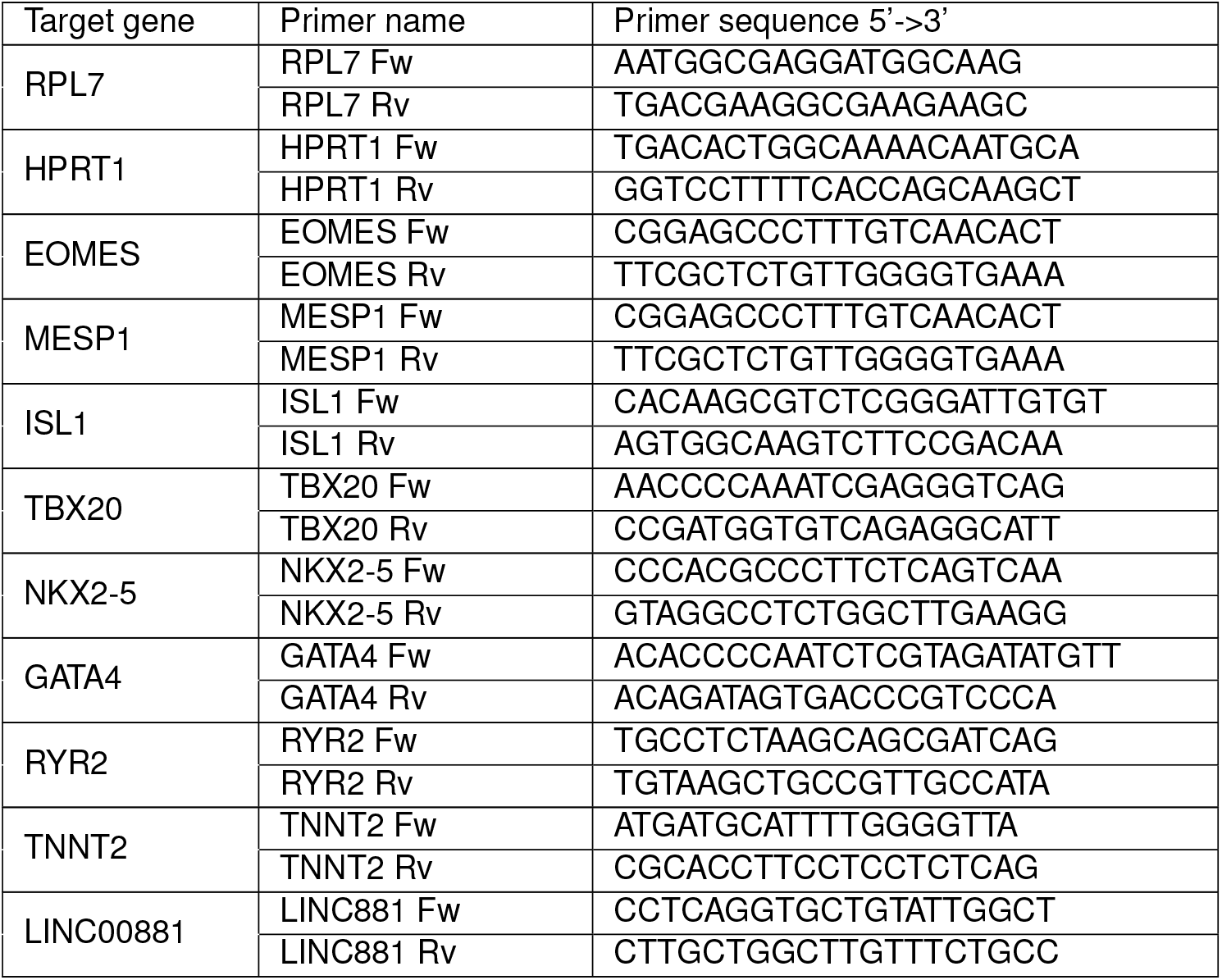
RT-qPCR primer sequences.

**Table 2.**
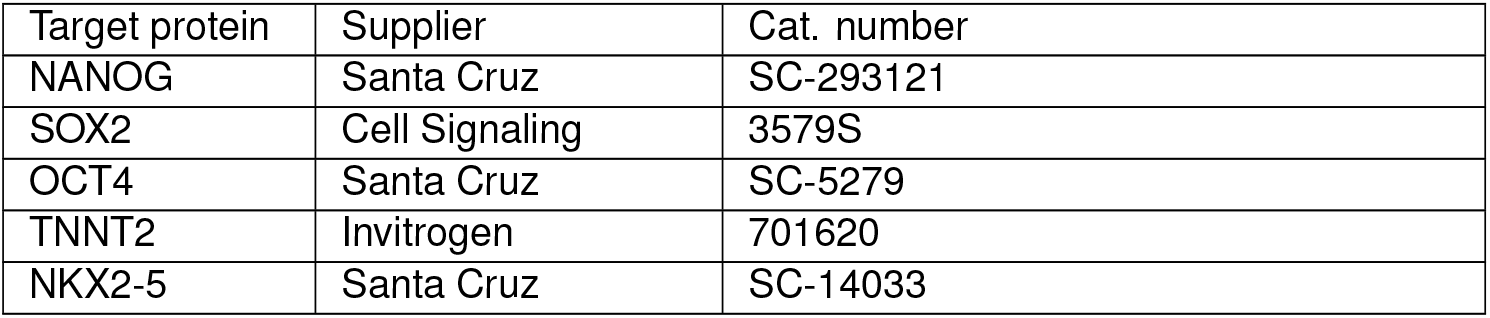
immunostaining primary antibodies.

| Target protein | Supplier | Cat. number |
| --- | --- | --- |
| NANOG | Santa Cruz | SC-293121 |
| SOX2 | Cell Signaling | 3579S |
| OCT4 | Santa Cruz | SC-5279 |
| TNNT2 | Invitrogen | 701620 |
| NKX2-5 | Santa Cruz | SC-14033 |

### Supplementary Figures

**Supplementary Figure 1:**
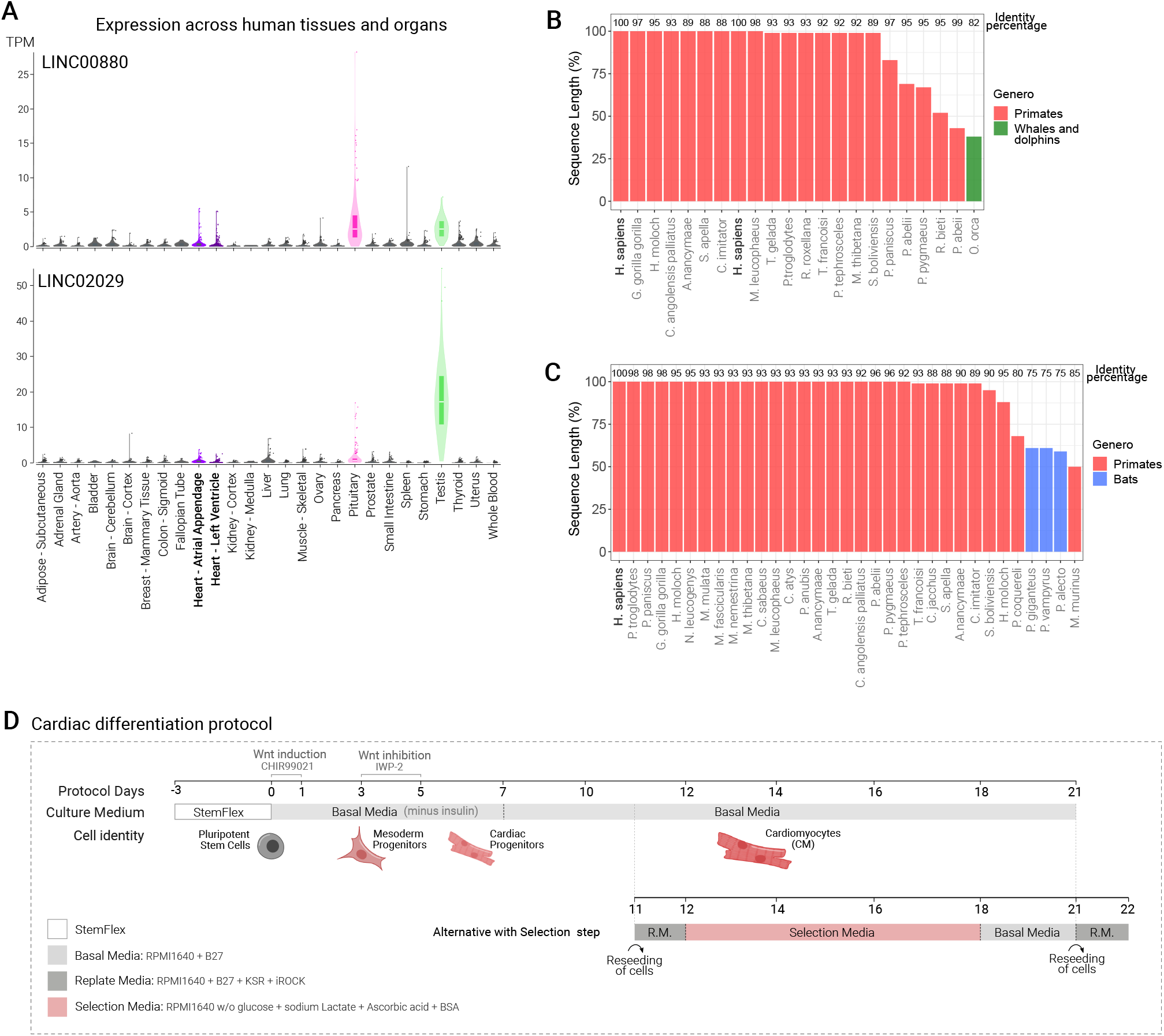
Characterization of neighbouring genes and species conservation of SHEN-LONG. A) Expression of LINC00880 and LINC02029 in human tissues and organs obtained from GTEx portal. Adult Heart tissue is shown in purple. Pituitary and testis are shown in pink and green respectively. B and C) Sequence alignment analysis using NCBI’s BLAST tool. Name of species is shown on x axis, lenght of sequence similarity is shown on the y axis as a percentage. Identity similarity is shown as a percentage on the top. D) Schematic of cardiac diffentiation protocol used. Alternative protocol with selection of cardiomyocytes is shown in the bottom.

**Supplementary Figure 2:**
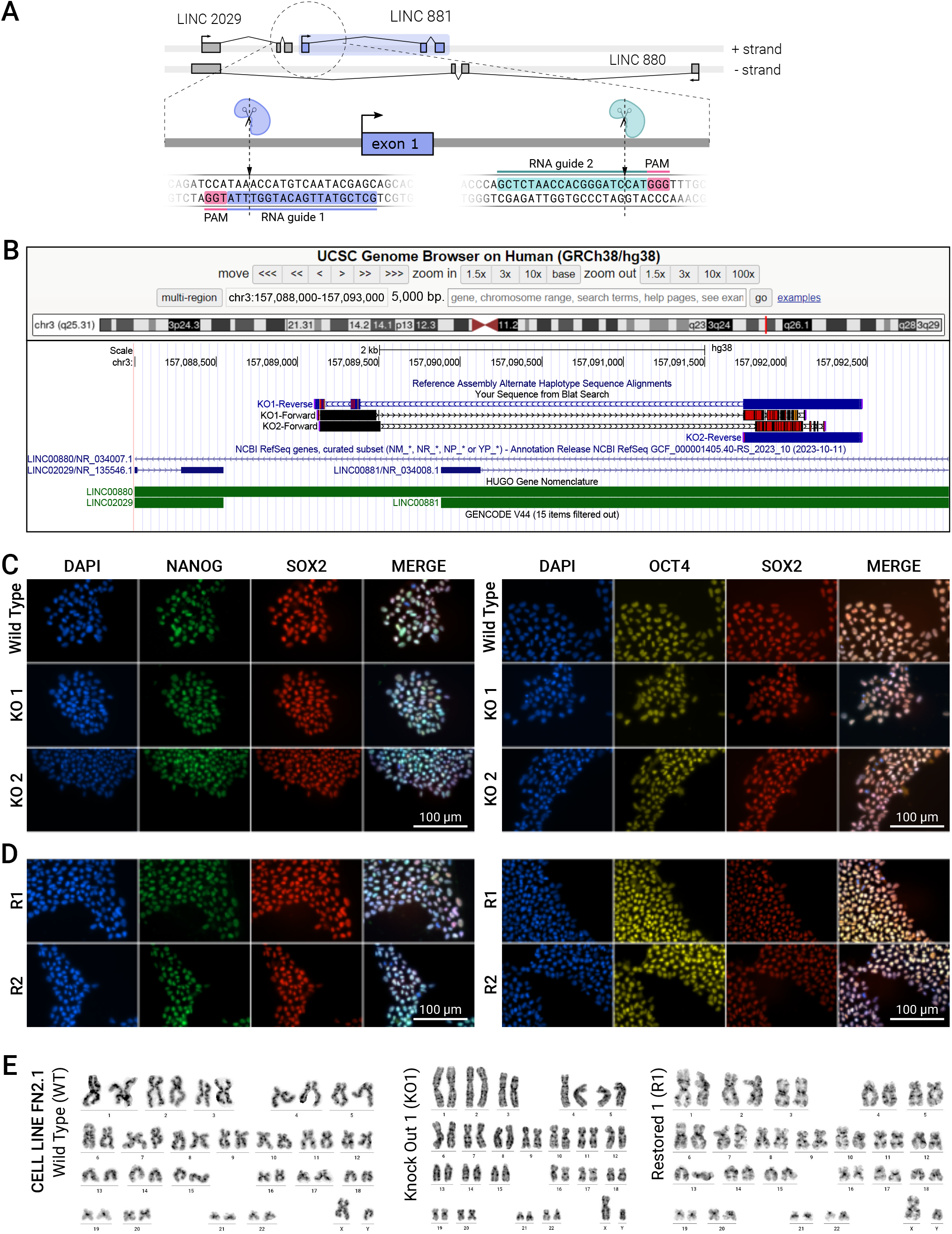
SHEN-LONG CRISPR Knock Out Validation. A) Schematic of CRISPR/Cas9 RNA guide relative position and sequence including PAM. B) Sanger sequence alignment of PCR amplicon corresponding to KO cell lines KO1 and KO2, showing deletion of target sequence. C) Immunofluorescence staining of pluripotency markers NANOG, SOX2 and OCT4 in WT, KO1, KO2, and D) R1 and R2 cell lines. E) Male normal karyotype (46,XY) of WT, KO1 and R1 cell lines.

**Supplementary Figure 3:**
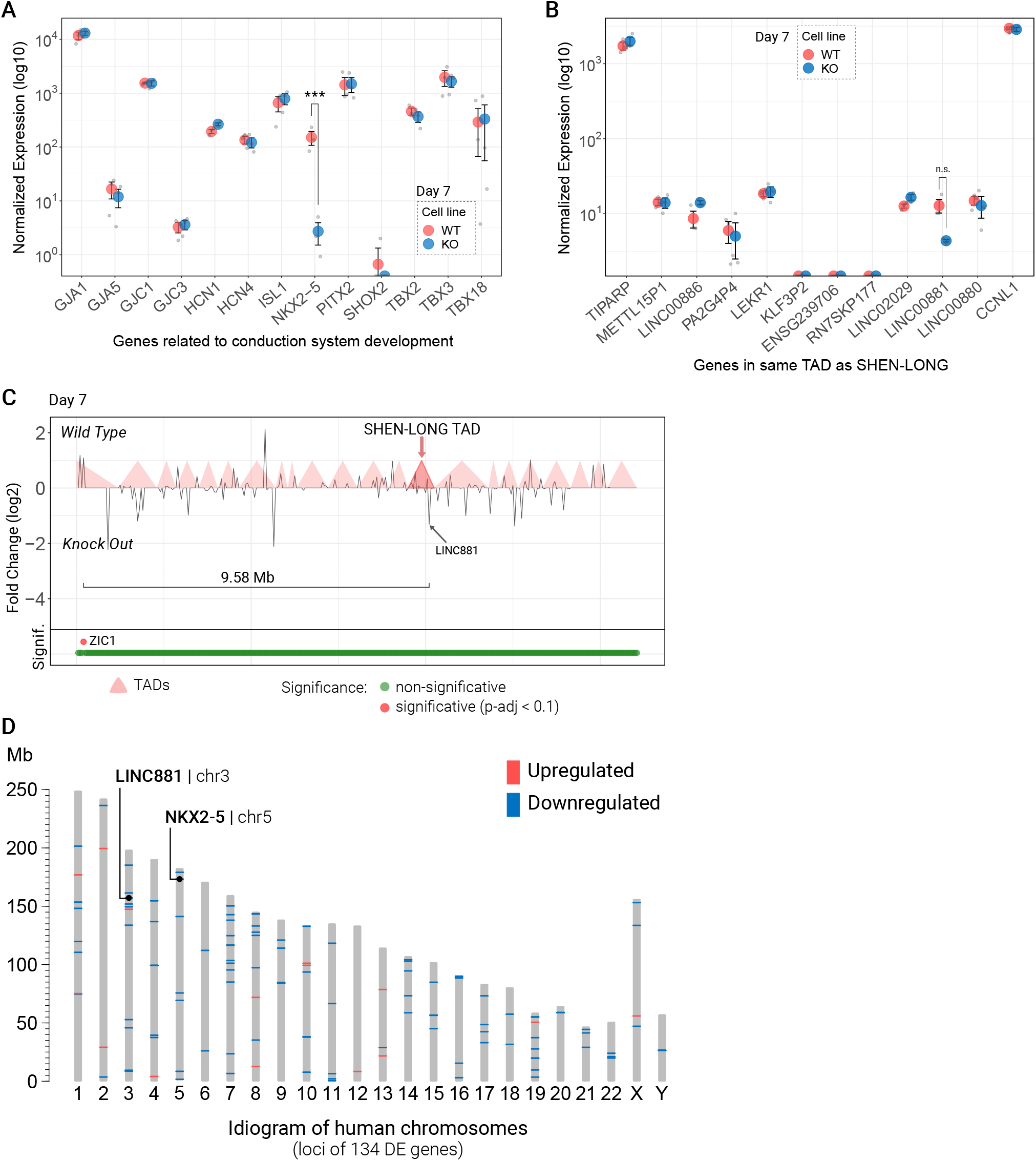
Differential expression analysis of SHEN-LONG KO and WT day 7 cardiomyocytes. A) Normalized expression levels in log10 scale of genes related to conduction system development. *** = p-adj*<*0.001 compared between KO (red) and WT (blue) cardiomyocytes at day 7. B) Normalized expression in log10 scale of genes localized in the same TAD as SHEN-LONG, compareing between KO (red) and WT (blue) cardiomyocytes at day 7. n.s: non significant. C) Relative localization, TAD and distance of diferentially expressed genes located within 10 Mb up and downstream of SHEN-LONG (distances not to scale) on day 7 cardiomyocytes. In total, 299 genes are shown. Statistical significance is shown in the bottom panel using p-adj*<*0.1 as significance threshold. The distance between the closest differentially expressed gene to SHEN-LONG is pointed out: ZIC1 at 9.58 Mb. TADs are shown as pale red rectangles. D) Idiogram of human chromosomes showing loci of all 134 differentially expressed between WT and KO cardiomyocytes at days 7 and 10. Upregulated genes are shown in red, downregulated genes are shown in blue. Localization of LINC881 on chromosome 3 and NKX2-5 in chromosome 5 are pointed out.

**Supplementary Figure 4:**
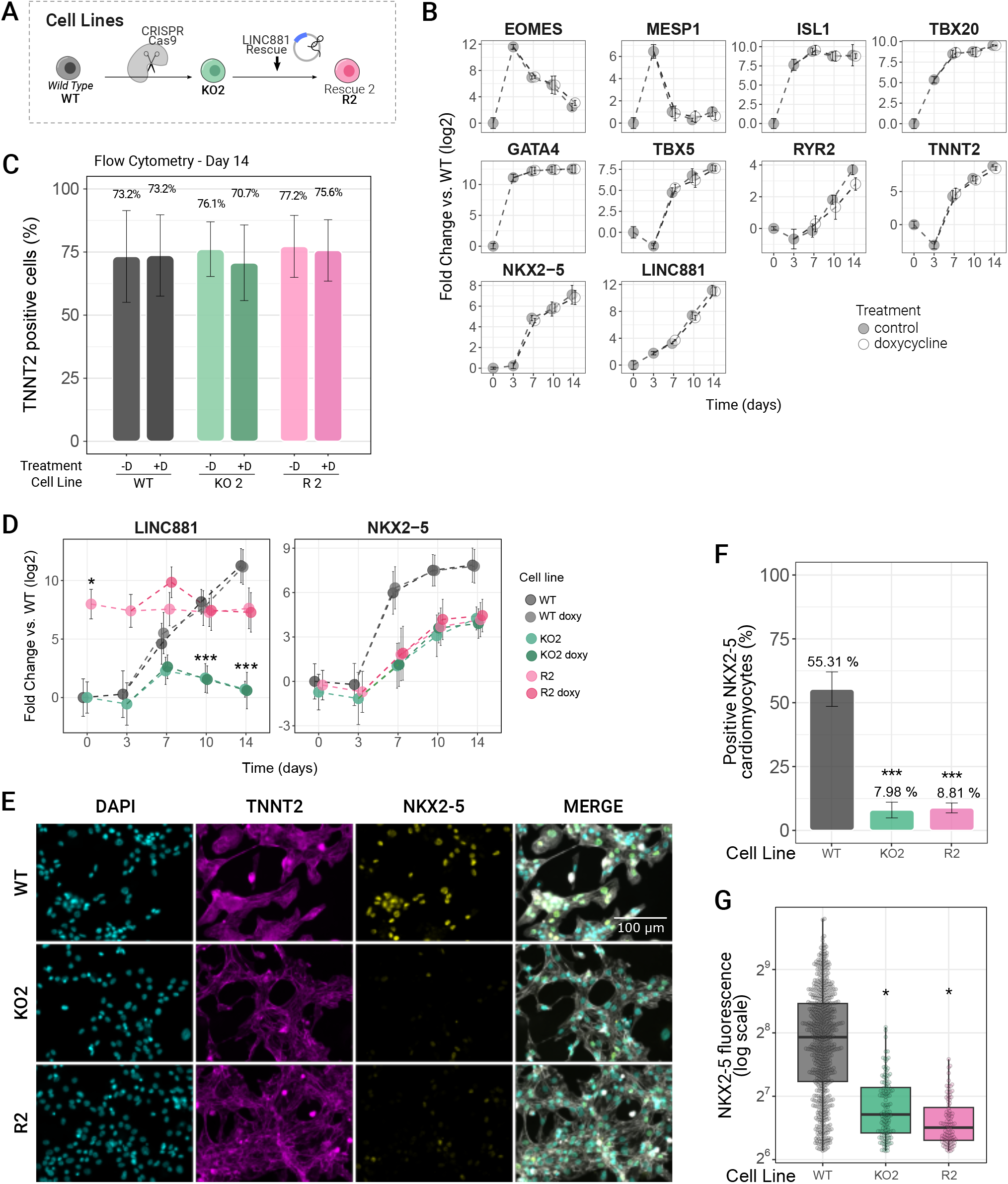
Cardiac differentiation of KO2 and R2 cell lines. A) Schematic of cell lines used B) Expression levels of cardiac differentiation markers in WT cells, with and without doxycycline treatment measured by qRT-PCR at days 0, 3, 7, 10 and 14: mesoderm commitment (*EOMEs* and *MESP1*); cardiac progenitor transcription factors (*ISL1, TBX5, TBX20, GATA4* and *NKX2-5*); cardiomyocyte proteins (*RYR2* and *TNNT2*); and cardiac lncRNA *LINC881*. Data is presented as mean of three biological replicates with bars as mean *±* s.e.m. C) Cardiomyocyte percentages at day 14 measured as TNNT2^+^ cells. D) Expression levels of LINC881 and NKX2-5 along cardiac differentiation protocol assesed by RT-qPCR in WT, KO2 and R2 cell lines. Data is presented as mean of three biological replicates with bars as mean *±* s.e.m. E) Immunofluorescence staining of NKX2-5 (yellow) and TNNT2 (magenta) at day 22 of cardiac differentiation. Nuclei were stained with DAPI (cyan). Merged channels are shown in the last column. F) Quantification of NKX2-5^+^ cardiomyocytes. Data is presented as mean of three biological replicates. G) Mean fluorescence of NKX2-5 per nuclei. Boxplot lines represent quartiles. Statistical analysis was performed with one-way ANOVA followed by Tukey test. Statistical significance: * = p*<*0.05, ** = p*<*0.01, *** = p*<*0.001.

**Supplementary Figure 5:**
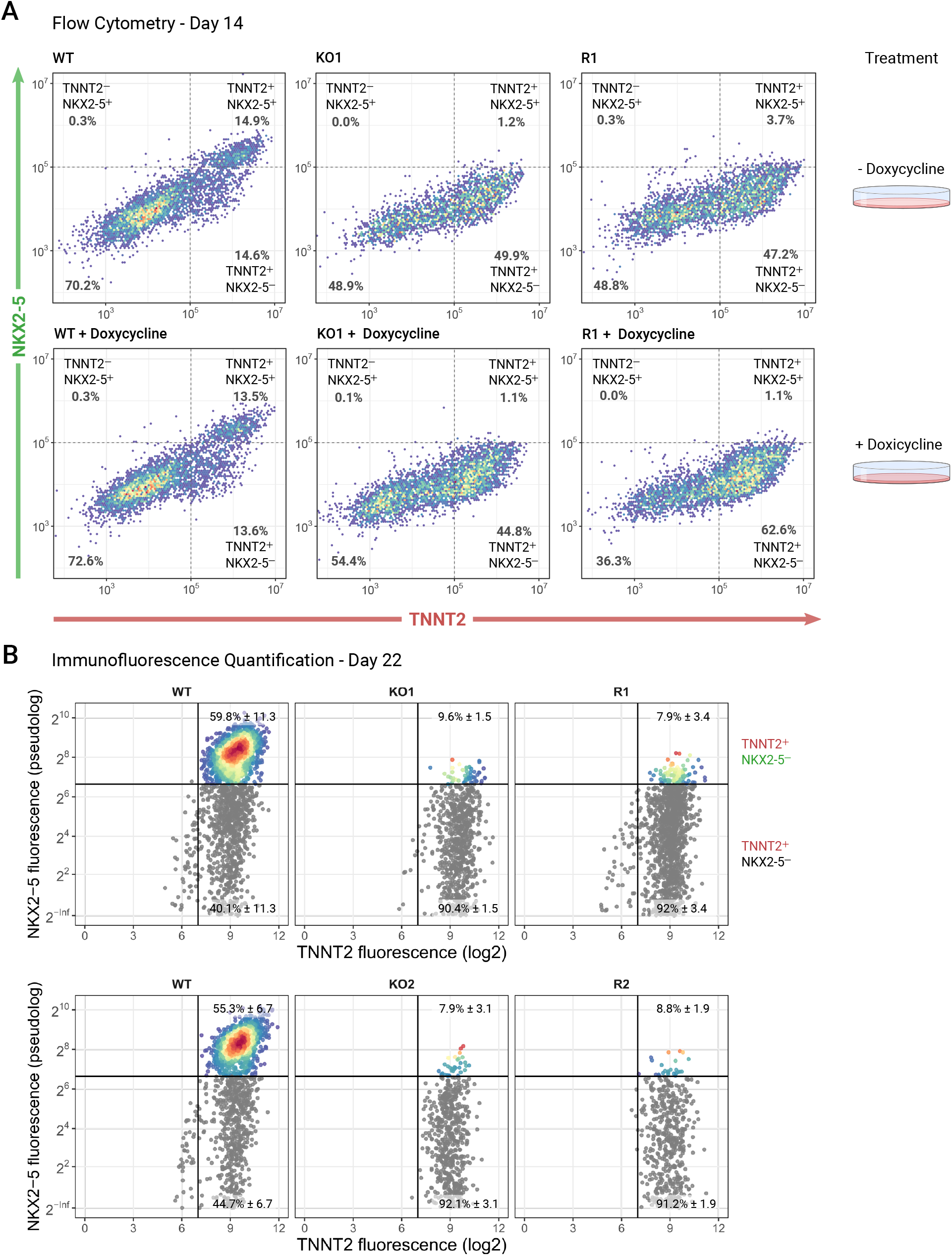
Analysis of NKX2-5 and TNNT2 protein expression in WT, KO and R cell lines. A) Representative flow cytometry of NKX2-5 and TNNT2 proteins in day 14 cardiomyocytes of WT, KO1 and R1 cell lines, with and without doxycycline treatment. B and D) Representative immunofluorescence quantification showing TNNT2+/NKX2-5+ cells in color. Percentages are shown for each condition. C and D) Percentages of positive NKX2-5 cardiomyocytes for each cell line.Data is presented as mean of three biological replicates with bars as mean *±* s.e.m. Statistical analysis was performed with one-way ANOVA followed by Tukey test. Statistical significance: * = p*<*0.05, ** = p*<*0.01, *** = p*<*0.001.

